# Rapid polygenic adaptation in a wild population of ash trees under a novel fungal epidemic

**DOI:** 10.1101/2022.08.01.502033

**Authors:** Carey L. Metheringham, William J. Plumb, William R. M. Flynn, Jonathan J. Stocks, Laura J. Kelly, Miguel Nemesio Gorriz, Stuart W. D. Grieve, Justin Moat, Emily R. Lines, Richard J. A. Buggs, Richard A. Nichols

## Abstract

Rapid evolution via small shifts in allele frequencies at thousands of loci are a long- standing neo-Darwinian prediction but are hard to characterize in the wild. European ash tree (*Fraxinus excelsior*) populations have recently come under strong selection by the invasive fungal pathogen *Hymenoscyphus fraxineus*. Using genomic prediction models based on field trial phenotypes and 7,985 loci, we show a shift in genomically estimated breeding values in an ancient woodland, between adult trees established before the epidemic started and juvenile trees established since. Using simulations, we estimate that natural selection has eliminated 31% of the juvenile population. Thus, we document a highly polygenic heritable micro-evolutionary adaptive change over a single generation in the wild.

**One-Sentence Summary:** Subtle changes at thousands of genomic locations in one generation allow woodland trees to respond to a new selective pressure

## Main Text

Whether complex traits typically adapt to new environments via large allele frequency changes in a few loci, or small allele frequency changes in many loci, is an open question in evolutionary biology (*1–5*). While theory suggests that a highly polygenic response should be rapid and effective (*3–6*), it is far easier in nature for population geneticists to demonstrate cases of natural selection involving low numbers of loci with large effect size (e.g. *7*, *8*–*10*). While the methods of quantitative genetics can show, by statistical comparison of related individuals, that additive genetic variance exists for complex traits under selection, it has often not been possible to show by these methods that response to selection is occurring (*11–14*). This situation has led to a disconnect between population genetics and quantitative genetics. Genomic prediction approaches, developed for agricultural breeding, that use genome-wide SNP data to predict individuals’ genetic merit for a quantitative trait of interest, can enable us to bridge this gap (*5*, *15*, *16*). If we can show genome-wide allele frequencies before and after the arrival of a new selective pressure (*17*) or in different age classes at a single time point (*12*, *18–22*), affecting genetic merit, a polygenic response to selection could be demonstrated.

The possibility of a rapid adaptive response is of particular interest in the case of the ash dieback epidemic that has swept across Europe in the past three decades, caused by the fungus *Hymenoscyphus fraxineus*, an invasive from East Asia (*23*). Numerous studies based on planted trials suggest that heritable variation in susceptibility to the fungus exists within European ash (*Fraxinus excelsior*) populations (*24–27*). The intensity of selection on viability may be stronger in smaller, younger ash trees, since they are observed to die more rapidly from ash dieback infection than larger, older trees (*28*, *29*). Rapid juvenile mortality may occur because the fungus can more quickly encircle the main stem and because smaller trees are closer to the leaf litter where *Hymenoscyphus fraxineus* apothecia are produced. Recruitment of the next generation may also be impacted by reduced reproduction by adult trees damaged but not yet killed by the fungus (*30*, *31*).

It has been hypothesized that mortality of susceptible juvenile trees and reduced reproduction by susceptible adult trees will drive changes in allele frequencies leading to an increase in disease resistance in the new generation of ash (*24*, *27*). In a previous study, based on 38,784 ash trees, ∼7 years old, from British, Irish and German provenances growing in field trials, we sequenced the 623 most healthy trees (score 7 on the scale of Pliura et al. (*32*)) and 627 trees whose woody tissues were highly damaged by ash dieback (scores 4 or 5). We used genome-wide association study to rank loci associated with these phenotypes by p-value (*26*) and used sets from 100 to 50,000 of the top loci to train genomic prediction models, which were tested on 148 trees. We found 10,000 loci to give Genomic Estimated Breeding Values (GEBVs) with the highest frequency of correct allocations of test trees (0.67) (*26*). We now calculate that these GEBVs explained 24.0% (CI: 11.5 - 37.0%) of the phenotypic variation in the test population’s damage due to the fungus (see Materials and Methods)

If natural selection is acting on natural ash populations under high disease pressure we would expect to see GEBVs increase in the younger generation of trees that have been exposed to infection since germination, with shifts in allele frequencies that correlate with their effect sizes. Here tested this hypothesis in a woodland located within the geographic sample range of Stocks et al. (*26*).

### Phenotypic and genomic characterisation of an ash population

Our study site, Marden Park wood, is an ancient semi-natural woodland dominated by *F. excelsior* where the pathogen *H. fraxineus* is thought to have been present since 2012 (Fig. S1). It is located in UK National Seed Zone number 405, a provenance included in the previous trials (*26*). Phenotypic assessments of this woodland in 2019, when we collected ash tissue samples for sequencing, found *H. fraxineus* symptoms on the majority of trees (Fig. 1), and no evidence for felling or removal of dead trees. Damage from the fungus had further increased by 2021 especially in juvenile trees (Fig. 1). This fits with widespread documentation of the ongoing progress of the ash dieback epidemic throughout Europe (*29*, *33*).

**Fig. 1.**
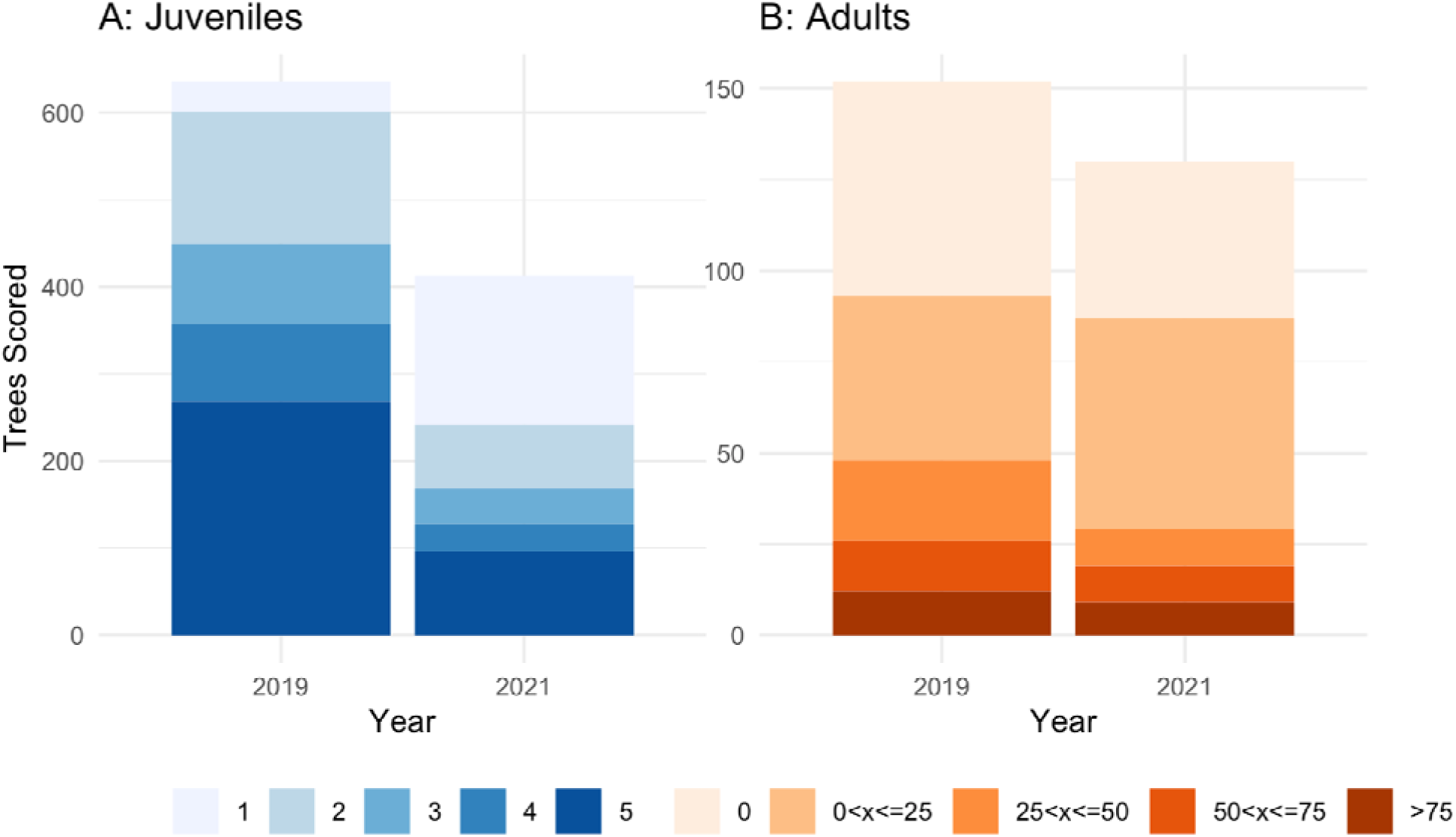
Phenotypic health assessments of trees that were labeled and genome sequenced in 2019. **(A**) Juvenile tree health scores (where 5 is the most healthy) in 2019 and 2021. (**B**) Adult tree percentage canopy coverage estimates in 2019 and 2021. A number of labels were missing by 2021, especially among juveniles, and only labeled trees were reassessed.

We generated short read sequence data for 580 individuals (128 adults, which had established pre-epidemic, and 452 post-epidemic juveniles; Fig. S2) and 30 technical replicates at approximately 11X whole genome coverage and estimated allele frequencies at ∼9 million SNP loci. Of the 10,000 SNPs used for genomic prediction in the trials (see above), 7,985 were variable in the Marden Park dataset and passed allele frequency and quality thresholds. This smaller number of polymorphic SNPs reflects the lower genetic diversity present in Marden Park wood than in the planted trials, which included many seed zones. Of the 2,015 SNPs that were not variable in the Marden Park population, 1,055 were fixed for the allele associated with low ash dieback damage in the planted trials and 960 for the allele associated with high damage.

We calculated GEBVs for the Marden Park trees from the 7,985 polymorphic SNPs using the parameters of the genomic prediction model trained on the field trials (*26*). Our visual assessments of ash dieback damage, scored on a five point scale for juveniles (a similar method to that used in the trials), and as percentage canopy cover in adults, showed no significant relationship with individual’s GEBV scores (Fig. S3 and 4). A weak relationship between individual GEBVs and phenotypes assessed in the field is expected due to the large environmental component of damage phenotypes in the wild, due to local microenvironments, age differences within cohorts and presence of other microorganisms. In addition, the phenotypic scoring method we used on adult trees is of necessity different to the method used to evaluate juveniles in the field trials. Hence, we might not expect the canopy cover of a mature tree in a woodland to be an accurate predictor of its breeding value for susceptibility to fungal damage in juvenile trees.

Instead of using the adult phenotype, breeding values are better estimated from the phenotypes of progeny when environmental effects are large (*34*). For 20 adult trees we were able to apply this approach by identifying offspring (n = 121) among the juveniles, using the sequoia R package (*35*) on 1000 SNPs. For these we found significant correlation between the GEBV of the parent trees and the mean health score of their offspring (r = 0.368, p = 0.029, n = 37, Fig. 2A), suggesting that our GEBVs are predicting breeding values relevant to the vulnerable juvenile stage (this will be an underestimate if selection has already eliminated the more susceptible individuals - see below). In contrast, our visual assessment of the parent trees’ health was a poor predictor of the mean phenotypic health score of the offspring (r = 0.112, p = 0.54, n = 37, Fig. 2B).

**Fig. 2.**
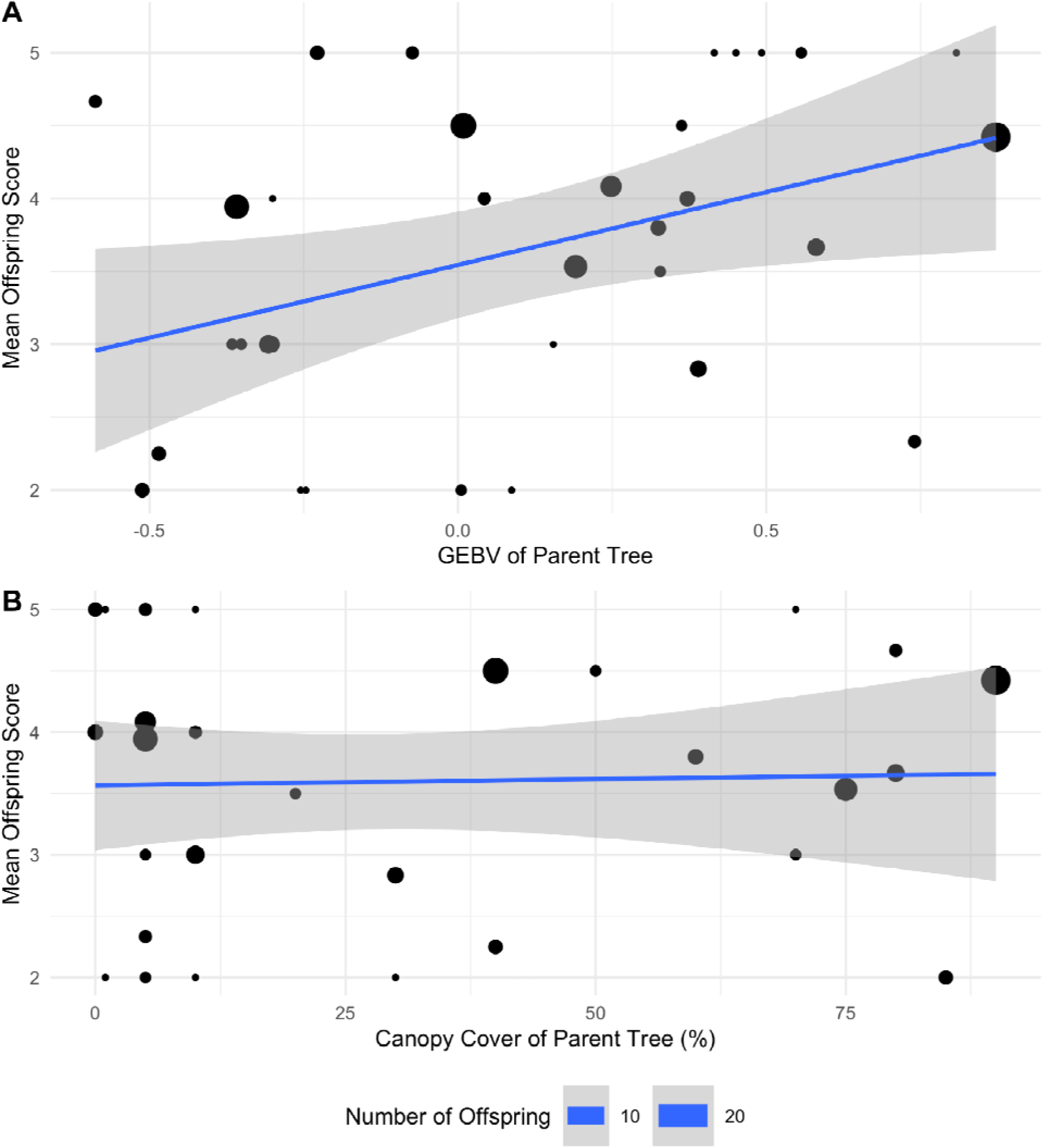
Correlation of (**A**) GEBV of parent trees (B) canopy cover of likely parent trees and with the mean phenotypic health score of juvenile offspring as assigned in sequoia, weighted by the number of offspring trees assigned to each parent.

A more systematic assessment of adult tree health was obtained from the rate of spring green-up from April to June in the 53 largest trees. These rates were estimated from three dimensional red-green-blue point cloud data using structure-from-motion analysis of sensor imagery from a consumer-grade uncrewed aerial vehicle. The progress of each tree’s greening was measured by the normalized green chromatic coordinate. Green-up occurred at a higher rate, *s*, in trees with higher GEBVs (Estimated *s*:GEBV interaction = 0.23, 95% CI = 0.01 - 0.53, Table S4, Fig. S5). This trend is consistent with the positive association between the degree of leaf bud burst in early May and resistance to ash dieback in a Lithuanian study (*36*).

### Evidence for allele shifts due to natural selection

We found a shift in mean disease susceptibility GEBV between adult trees and juvenile trees in our population. The mean GEBV of juvenile trees (μ = 0.22, n = 452) was higher than the mean GEBV of adult trees (μ= 0.15, n = 128; Fig. 3A), with a shift of 0.07 (95% confidence intervals: −0.016, 0.142). The statistical significance of this increase in score cannot be evaluated by treating each juvenile tree as an independent observation, since some adults have left multiple offspring. Instead we compared the GEBV score of each juvenile individual with that predicted from its ancestry. If the surviving juveniles have atypically high GEBV scores, then on average their scores should be higher than predicted from their ancestry. To capture this ancestry, we characterized relatedness between juveniles and the adults by selecting 5,793 loci, which do not contribute to our estimate of breeding value, that were not in close linkage with the 7,985 GEBV sites or with one another, had no missing data and had minor allele frequencies of over 0.3. The additive relatedness matrix among adults and juveniles calculated from these 5,793 loci was used to predict the juvenile GEBVs from the actual GEBVs of related adults. These predictions were significantly lower (p = 0.001, t-test of the intercept in regression) than the actual juvenile GEBVs (regression in Fig. 3B), giving a shift of 0.054. The higher than predicted GEBV in the living juveniles would be expected if part of their cohort with lower GEBV had not survived. This difference is smaller in magnitude than the 0.07 difference between adult and juvenile means. We would expect the 0.054 value to be an under-estimate, since it deliberately excludes the variation in mating success of adults, since those effects could include random differences in fertility and fecundity which are components of genetic drift (not selection).

**Fig. 3.**
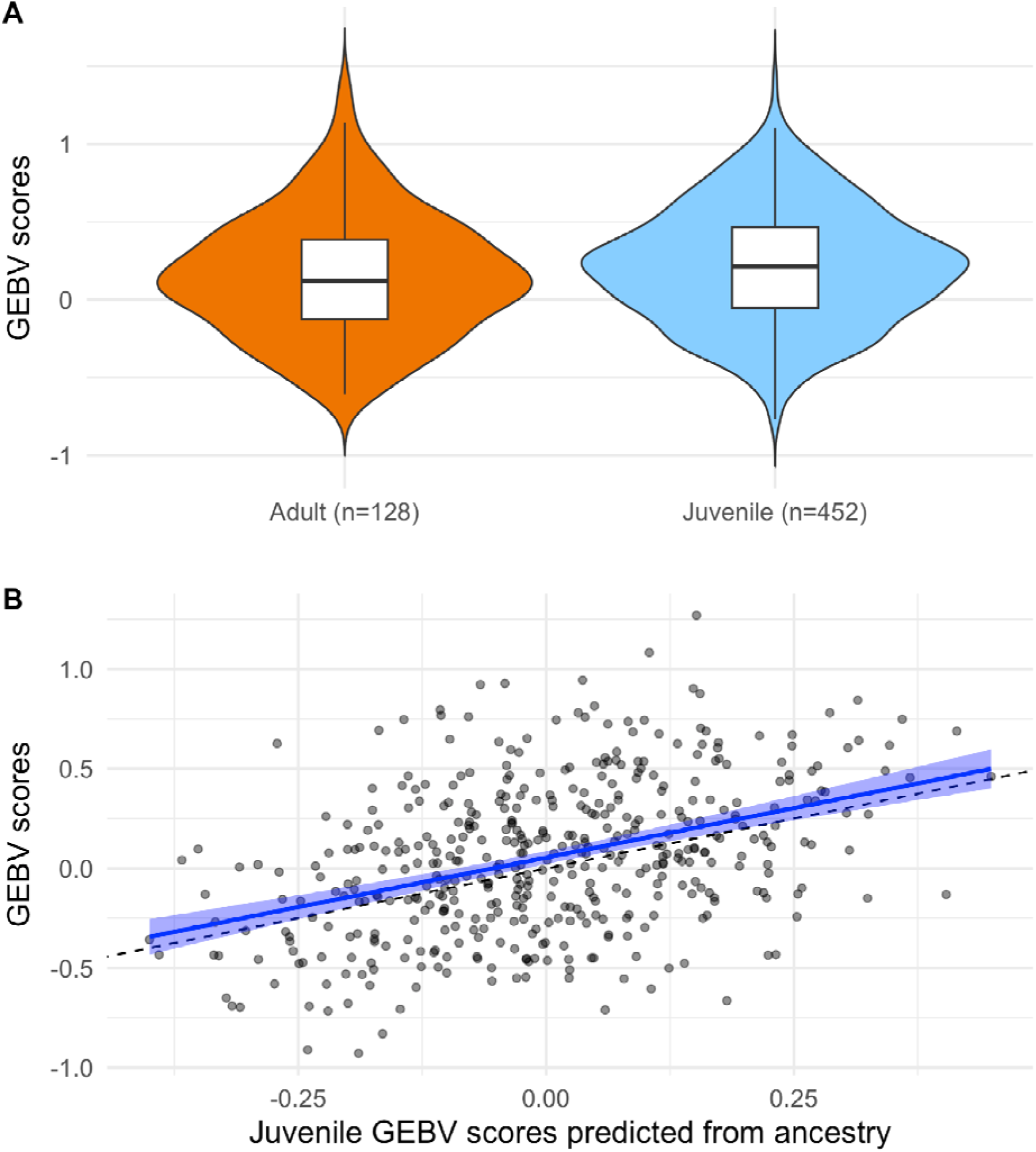
Shift in GEBV between adult and juvenile trees. (**A**) Genomic estimates of breeding values for 128 adult trees and 452 juvenile trees calculated from 7,985 SNP loci, using the model trained on large planted trials. (**B**) Each juvenile tree’s GEBV based on the 7,985 loci, plotted against an alternative prediction based on ancestry alone via an rrBLUP model trained on the adult trees’ GEBVs. The fitted regression line (shown in blue) is significantly greater than the 1:1 line (shown in dashed black), demonstrating higher than expected GEBVs in the juvenile cohort. Standard error of the fitted regression is shown by the blue highlighted area.

Three additional lines of evidence show that the changes in allele frequencies between parents and offspring are due to selection.

First, the variance in allele frequency between parents and juveniles are much greater at GEBV loci than at unlinked putatively neutral loci, unless we remove the effects of selection in the former. The inter-generational change for the 5,793 neutral loci was F = 0.00033 (SE: 0.00005) indicating minimal genetic change between adult and juvenile cohorts, corresponding to an effective population size, 1/2F, of 1,515. In contrast, the 7,985 GEBV loci had over double the variance of F = 0.00079 (SE: 0.0007), which is significantly greater (P = 0.0006, permutation test). To estimate the effect of drift on the GEBV loci, we modelled and removed the selective component of change, modelled as a function of effect sizes and frequency of alleles at each locus, yielding a drift estimate of F = 0.00028, a value that is very close to, and not significantly different (P = 0.65) from the value for the neutral loci. Thus, the contribution of genetic drift to changes at the GEBV loci is indistinguishable from the neutral loci, and the larger allele shift in the GEBV loci is attributable to selection.

Second, we also detect a pattern whereby the direction and size of changes are proportional to the estimated effect sizes and frequency of alleles at each locus, so that larger changes are at alleles at intermediate frequencies with large effect sizes. (Fig. 4). The regression was highly significant (r = 0.0015, p = 0.00038, F = 12.64 on 1 and 7983 DF). This pattern also supports the case that the changes at GEBV loci are caused by selection rather than chance. To test whether the regression was affected by the precision of the estimates of *E_i_* for each locus (as measured by its standard error *SE_i_*) the regression was repeated with each locus inversely weighed by 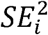. The result was essentially unaffected, with the regression coefficients differing by only 0.03%.

**Fig. 4.**
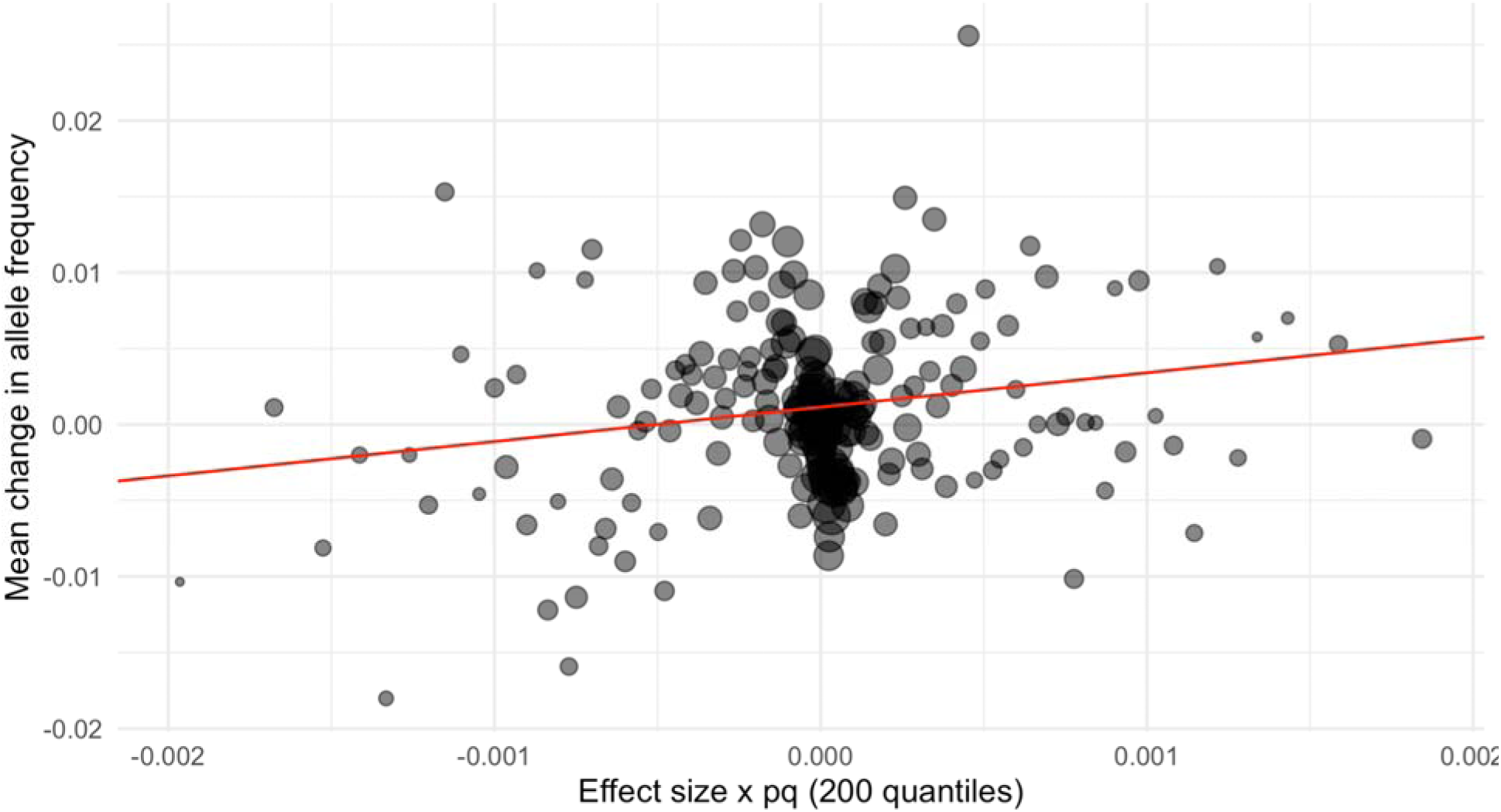
Mean change in allele frequency between adults and juveniles. Mean change in allele frequency plotted against the product of effect size and minor allele frequency. The values for 7,985 sites have been combined into 200 points. Each point represents the average of 39 or 40 sites (adjacent values of Effect size x pq, binned), with area representing the precision of the mean change (1/var(Δ*f*)). The red line is the fitted linear regression.

Third, an individual-focussed prediction of the change in GEBV allele frequencies can be directly tested in the case of the 121 juveniles for which we were able to assign at least one parent with confidence (see above). In these cases we could predict the allele frequencies expected by random transmission. The observed frequencies were significantly different from these random expectations (p < 0.05, Wilcoxon rank sum test with Bonferroni correction). The advantageous alleles, associated with low susceptibility to ash dieback, were over-represented, the excess frequency being correlated with the effect size (r = 0.0595, p < 10^-6^). This pattern again suggests that the offspring carrying these alleles had a selective advantage, making them more likely to survive to become part of our sample (compared to siblings that did not).

To further explore the polygenic nature of the shift in GEBVs between adults and juveniles in Marden Park, we calculated the proportion of total variance in GEBVs contributed by the loci on each 1 kbp segment of our contig assembly. To explain 90% of the variance required 2,123 of these segments, which carried 3,752 GEBV SNPs (out of 7,985), which are on 1,342 different contigs (out of 2,783 carrying GEBV SNPs). In order to exclude the possibility that the shift in GEBV between parents and offspring was explained by a few segments of large effect, we re-ran the analysis excluding the segments of the genome accounting for the top 25% of variance. The remaining loci still showed a substantial and highly significant shift of 0.046 (p = 0.0012) between juveniles’ GEBVs and that predicted for them by their relatedness to adults. This consistent pattern across the genome underlines the highly polygenic nature of the selection occuring in the woodland.

### Simulation

To evaluate the effectiveness of our study design in estimating small effects at such a large number of loci, and to quantify selection over a single generation, we conducted a simulation of our entire study. First, we simulated the pooled-GWAS and genomic prediction experimental design (*26*). Genotypes at 7,985 loci were generated as random binomial draws using the empirical estimates of the allele frequencies. Susceptibility phenotypes were calculated for these genotypes using the empirical estimates of the corresponding effect sizes and a random environmental effect that conferred a heritability of 0.4. High and low susceptibility pools were generated corresponding to the proportion of trees selected for the pools in the real experiment. These pools were used to estimate a relative effect size for each locus using rrBLUP. The relative effect sizes were estimated convincingly (Fig. S6), albeit with some error (r^2^=0.30). When the effects at multiple loci are summed, they allow for accurate genomic prediction (r^2^=0.39, Fig. S7).

Second, we simulated a population with the observed adult GEBV scores and the observed additive relationship matrix at the neutral loci. We implemented selective deaths caused by a range of truncation selection intensities on the underlying latent variable (the score describing susceptibility to the fungus). The implementation allowed for the reduced response to selection due to expected error on our estimate of effect sizes influencing the realized heritability (h = 0.24). By regressing the shift in GEBV score on the quantile of the truncation selection, we were able to estimate that the observed shift of 0.054 would have been produced by the selective elimination of the lower 31% of the latent variable scores.

The magnitude and direction of the change in allele frequencies between generations (Δ*p*) in the simulation were comparable to those found in our field study. There was a highly significant regression of Δ*p* on *<u>p</u>*(1 - *<u>p</u>*)*E*, with a relatively low correlation (multiple r^2^ = 0.0011, p = 0.0025, F = 9.102 on 1 and 7983 DF; Fig. S8). The small r^2^ value is explained by the relatively large error in estimates of E at each locus combined with stochastic variation in the allele frequency change; whereas the combined evidence over many loci is highly significant.

### Conclusion

We have thus documented a heritable, micro-evolutionary adaptive change occurring over a single generation due to small allelic shifts in thousands of loci. To have demonstrated a shift in additive genetic variation without genome-sequencing would have involved producing large numbers of clones or offspring from many adult and juvenile trees in the woodland, and growing these in uniform conditions for several years under ash dieback inoculum pressure. Even then, such an approach would not have given us information about the number of loci involved in the adaptive shift. We were only able to characterize this highly polygenic shift in allele frequencies because we could use estimates of effect sizes at genome-wide loci obtained from large field trials. Our approach also relied upon the existence of multiple generations co-existing in nature, the older of which is known to die more slowly, and so retains the ancestral allele frequencies. It also required population genome sequencing of a large number of individuals, which was facilitated by the relatively small size of the ash genome (∼880 Mbp).

Few other studies have thus far sought evidence for micro-evolutionary change via shifts in GEBVs (*15*), though several studies in animals have examined shifts in predicted breeding values based on pedigrees (*37*). A recent study showed an increase in GEBVs calculated using genomic prediction algorithms for adult weight in a 35 year data set for Soay sheep which aligned with changes in predicted breeding values (from pedigree based studies), though phenotypic body weight decreased over this time span (*16*). In the present study, by contrast, we used the difference between predicted GEBV (based on relatedness) and observed GEBV (based on loci associated with the trait of interest) to infer that changes in allele frequency could be attributed to selection. These strategies could be extended to other organisms in the search for highly polygenic allelic shifts associated with a trait under selection, to produce neo-Darwinian evolution. It is likely that such shifts are common (*5*). An analogous approach in humans could be to use comparison of polygenic risk scores in ancient and modern DNA.

The action of natural selection in the wild for increased health of European ash under ash dieback is good news. Selection for low ash dieback susceptibility may occur partly by reduced seed or pollen production from adult trees damaged by ash dieback (*30*, *31*, *38*) resulting in non-random mating within the adult population and partly through rapid death of young trees infected by ash dieback, constituting a missing fraction that did not survive. The significant shift which we detected in the GEBVs of juveniles compared to their GEBVs based only on relatedness to the adults must by definition be demonstrating the latter selective mechanism. Whether the rate of change we observed will be sufficient for evolutionary rescue to occur (*39*), and whether its end point will be a fully resistant tree, is as yet unknown. The population we studied is fixed for alleles associated with high susceptibility at more than 900 loci; at these loci gene flow from other populations will be essential to introduce beneficial alleles. Our parentage analyses suggest ongoing migration of pollen and/or seed into the population, but this may decrease as ash becomes more scarce in the landscape. As low susceptibility is highly polygenic, the speed of evolution may be inhibited by linkage among loci (*40*, *41*). As the epidemic progresses, selective pressure may decrease as the pathogen follows the host demographics to become more scarce in the landscape. This will diminish the speed of evolution, especially given that heritability may be low in highly variable woodland environments. In many British ash woodlands, natural regeneration of ash is scarce due to deer browsing, so deer management will be essential for natural selection by *H. fraxineus* to be effective. Further interventions could enhance the rate of evolutionary change, for example, by introducing genetic diversity from other seed sources or from trees selected, or bred, for high GEBV.

Several breeding programs for ash dieback resistance are currently underway in Europe, with parental trees selected based on their ash dieback damage phenotypes. Our results suggest that a more effective approach may be to select parental trees based on their GEBV. A breeding program for increased disease resistance based on GEBVs would allow more accurate selection of parents with high breeding values, and would also allow parental pairs with complementary sets of alleles associated with resistance to be selected for crossing.

## Acknowledgments

We thank the Woodland Trust for access to Marden Park wood, and Tim Hodges and Kate Harvey for their support and assistance. This research utilized Queen Mary’s Apocrita HPC facility, supported by QMUL Research-IT. http://doi.org/10.5281/zenodo.438045

## Funding

UK Government Department for Environment Food and Rural Affairs (Defra) grants under the Future Proofing Plant Health Programme and the Centre for Forest Protection to RJAB

Department of Agriculture, Food and the Marine, Ireland, Teagasc Walsh Fellowship to WJP

Natural Environment Research Council London DTP PhD studentship to WRMF.

UKRI Future Leaders Fellowship (MR/T019832/1) to ERL.

## Author contributions

Conceptualization: CLM, RJAB, RAN

Methodology: RAN, CLM, WRMF, RJAB, JM

Fieldwork: CLM, WJP, WRMF, JJS, LJK, MNG, RAN, RJAB, SWDG, ERL

Investigation: CLM, RJAB, RAN, WRMF

Visualization: CLM, JM, RAN

Funding acquisition: RJAB

Project administration: RJAB

Supervision: RAN, RJAB, ERL, SWDG

Writing – original draft: CLM, RJAB, RAN

Writing – review & editing: CLM, WRMF, RJAB, RAN

## Data and materials availability

Raw reads for all sequenced samples are available on the ENA short read archive under accession PRJEB44697 (ERP128769). Analysis scripts and associated data are available in the Zenodo repository https://doi.org/10.5281/zenodo.10808942 and are available on Github at https://github.com/CareyMetheringham/MardenPark..

## Supplementary Materials for

### Materials and Methods

#### Site Selection

For this study we required an accessible ash-dominated woodland site in the south east of England, where *Hymenoscyphus fraxineus* has been present since 2012 (the year when the fungus was first found in England). We sought a site with the following characteristics: containing ash trees of a wide range of ages and sizes, including the presence of young naturally regenerated saplings; the older ash trees should be (as far as could be ascertained) naturally seeded rather than planted; the owners should have conducted few management interventions since 2012, and herbivore populations should be low enough to allow ongoing natural regeneration. We examined potential sites that fit these criteria, with Woodland Trust and Kent County Council. In August 2019 we selected a site within Stubbs Copse in Marden Park Wood in Surrey (51.2707, −0.0400) that has belonged to the Woodland Trust since 1994. This is on a chalk plateau of the North Downs at approximately 244 m above sea level. It is in the Surrey Hills Area of Outstanding Natural Beauty and is a Site of Special Scientific Interest. It is an ancient semi-natural woodland site where mature and pole stage ash is dominant; with beech, whitebeam and oak as rarer components. It approximates to National Vegetation Classification (NVC) W12a - beech/oak/ash with a dog’s mercury sub-community. When the Woodland Trust surveyed the site in 2019 they found that ash dieback was prevalent and only a “handful” of trees showed no signs of the disease with others showing “advanced stages of decline, especially in the younger, pole-staged trees”. The 50 years plus management objectives for the wood are to increase its resilience to pests and diseases and maximise its biodiversity. The only management interventions planned from 2019-2024 were the removal of non-native species and the coppicing of some hazel in the understorey. Ash were only to be felled where they pose a risk to site visitors and footpaths. The Marden Park population is located in UK National Seed Zone number 405. Trees from this seed zone were present as one of the provenances used for training the genomic prediction model by Stocks *et al.* (*26*).

#### Sample and Phenotype Collection

From 16-20 September 2019 we surveyed and collected tissue samples from 784 ash trees in Stubbs Copse. We sought to exhaustively sample all ash trees, moving outwards from a central point. For each ash tree we recorded its size, level of ash dieback damage, and GPS coordinates. We labeled each tree with a plastic tag. We took a leaf sample from each tree, and when leaves were unavailable or inaccessible we took buds or twigs. Failing the availability or accessibility of any of these tissues, we took a cambial sample from the trunk using a hammer and chisel, cutting out an area of bark approximately 2 cm x 8 cm. Sampled tissues were immediately placed into plastic bags containing silica gel, to dry. For trees less than approximately breast height, we measured tree size as height from ground level, and we scored ash dieback damage using a scale similar to that of Pliura (*32*) with the difference that we had fewer categories of dead trees. Trees we scored as 1 were completely dead, 2 were mostly dead but with small sections of the main stem still living, 3 had infection which had spread into the main stem, 4 had minor infections and 5 were healthy trees with no signs of infection in the stem. For larger trees we measured size as diameter at breast height (DBH) and scored ash dieback damage as an estimate of percentage leaf canopy loss from visual inspection from the ground. On 16 October 2019 and 21 and 22 January 2020 we re-collected tissue samples from trees where the first DNA extraction attempt had failed, and as far as possible added aluminium tags to the trees included in the study. On 4 and 12 March 2021 positions of trees were collected using an RTK GPS (Bad Elf Pro), using NTRIP (Networked Transport of RTCM via Internet Protocol) connection to the Imperial University unit (LICCOOGBRO), approximately 27 km from the site. On 18 and 19 August 2021, 126 of the adult trees and 355 of the juvenile trees were re-phenotyped, and their locations mapped in detail (Fig. S1). The phenotypes showed damage due to *H. fraxineus* in juvenile trees increased between 2019 and 2021, with the mean health score decreasing from 3.6 (SE = 0.1, n = 642) in 2019 to 2.5 (SE = 0.1, n = 418) in 2021 (Fig. 1).The percentage of juvenile trees observed to be dead increased from 5.5% in 2019 to 40.9% in 2021. For adult trees, mean percentage canopy coverage was 22% (SE = 2.9, n = 160) in 2019 and 20% (SE = 2.3, n = 126) in 2021 (Fig. 1). The lower number of trees re-phenotyped in 2021 was most likely due to loss of labels and accidental loss of juveniles due to trampling or browsing. The 224 juvenile trees that were missing in 2021 had the same average health score (3.6, SE = 0.1) in 2019 as all juvenile trees scored in 2019. Thus it seems unlikely that their disappearance was mainly due to mortality caused by ash dieback. Moran’s I (Moran, 1950) was calculated in order to estimate the degree of spatial auto-correlation within the phenotypic data of trees which had their positions collected in 2021. The 2019 health scores of the juvenile trees which were mapped in 2021 showed significant spatial auto-correlation (Moran’s I = 0.127, p = 0) but this was not the case in 2021 (Moran’s I = 0.0135, p = 0.327). The health of the adult trees did not show significant spatial auto-correlation in either year (2019: Moran’s I = 0.0546, p = 0.0538; 2021: Moran’s I = 0.0452, p = 0.137). Spatial autocorrelation among juvenile trees may be a product of local biotic and abiotic factors, or of relatedness between juveniles in close proximity.

#### UAV phenotyping

We measured the greenness of leaf canopy for large adult trees within our sample site, in each month from February to June 2021, using a DJI Mavic Mini UAV (DJI, Shenzhen, China) with a 12 MP RGB sensor. Flights were conducted at 50 m and 70 m altitudes with both −90 and −55 degree camera angles, referenced to the horizon. Camera image acquisition parameters were set to “auto”. Images had a front overlap of 95% and a side overlap of 80% and were captured over an area 3.2 ha centered on the 0.6 ha study plot. Nine checkerboard ground control points were placed around and within the plot and precisely located using a Leica Total Station (Leica Geosystems Limited, Milton Keynes, UK). Structure from motion point clouds were constructed using Agisoft Metashape version 1.7.4 (Agisoft LLC, St. Petersburg, Russia) filtered using the Point Cloud Library statistical outlier removal and Euclidean cluster filters and downsampled using a point-to-point minimum distance of 0.05 m.

We also scanned the 0.6ha plot with a Riegl VZ400i terrestrial laser scanner (RIEGL Laser Measurement Systems GmbH, Horn, Austria) in May 2021, with 58 scan locations and an upright and horizontal scan in each location. Point clouds were filtered and downsampled in the same way as the UAV data. Individual tree crowns were semi-automatically segmented using the Forest Structural Complexity Tool (*42*) and manually refined. The TLS and UAV point clouds were co-registered using the ground control locations captured by both sensors. Trees were labeled using a co-registered stem map generated using a Leica Total Station. In total 120 trees were delineated. Individual trees were segmented in the structure from motion point clouds using the Open3D Library in Python (*43*), with a point neighborhood approach based on the semi-automatically segmented TLS point clouds. Within each tree the trunk was segmented from the crown using the width of horizontal slices. Crown greenness was calculated as green chromatic coordinate (gcc): g_mean_/ (r_mean_+g_mean_+ b_mean_).

#### DNA extraction and sequencing

DNA was extracted from dried plant material using Qiagen DNeasy Plant Pro Kits or an adapted CTAB extraction procedure (*44*). DNA for 755 trees, including 52 replicates was sent to Novogene for sequencing. Library preparation with NEBNext DNA Library Prep Kit and paired end Illumina sequencing on NovaSeq 6000 was carried out by Novogene (Germany) with a total amount of 1.0 g DNA per sample. Genomic DNA was randomly fragmented to a size of 350bp by shearing, then DNA fragments were end polished, A-tailed, and ligated with the NEBNext adapter for Illumina sequencing, and further PCR enriched by P5 and indexed P7 oligos. PCR products were purified using the AMPure XP system and resulting libraries were analyzed for size distribution by Agilent 2100 Bioanalyzer, quantified using real-time PCR, then sequenced.

#### Sequence analysis

We generated 0.71 to 19.92 Gb of read data per tree (μ = 9.93 Gb), giving 0.81 - 22.64 X coverage (μ = 11.28) of the whole genome (assuming a genome size of 877Mbp (*45*)) for 649 individuals and 30 technical replicates. Reads containing adapters, more than 10% undetermined bases or with more than 50% low quality (Qscore<= 5) bases were filtered out with SAMtools (V1.9)(*46*) and remaining reads were aligned against the BATG0.5 *F. excelsior* genome using the bwa mem function in the Burrows Wheeler Aligner (V0.7.17)(*47*) with minimum seed length of 19, matching score of 1, mismatch penalty of 4, gap open penalty of 6 and gap extension penalty of 1. For each sample, 3.0% to 98.9% of reads mapped to the *F. excelsior* reference genome (μ = 86.3%). Samples with less than 50% of reads aligning to the reference genome were excluded from further analysis.

SNP calling was performed using HaplotypeCaller, GenomicsDBImport and GenotypeGVCFs in the Genome Analysis Toolkit (GATK V4.1.4.0)(*48*). Sites with greater than 25% missing data were removed and hard filtering was performed on remaining SNP variants using VariantFiltration in GATK (V4.2.1.0) with the following parameters; read depth ≥ 10 and <200, variant confidence ≥ 20, QD (variant confidence normalized by read depth) ≥ 4, mapping quality ≥ 40, FS ≤ 60 and minor allele frequency ≥ 0.01. After excluding samples with low alignment and filtering our read alignments for quality, coverage and indels, we called SNPs at 28,499,977 variable sites in 600 samples, including 25 replicates. Linkage disequilibrium pruning was performed using PLINK 1.9 (*49*) with an independent pairwise r^2^ threshold of 0.5, window size of 50 and step size of 5 using the option --indep-pairwise, resulting in a reduced set of 4,287,210 SNPs.

In a PCA of the SNP data performed in PLINK, the first and second principal components explained 12.1% and 9.2% of the genetic variance respectively (Fig. S2). The identity of replicate samples was confirmed by calculating the genomic distance between samples using the –distance function in PLINK, with genomic distance recorded as 1 minus identity-by-state. The 25 technical replicate pairs had genomic distances of between 0.051 and 0.094 (μ = 0.063) (Table S1) whereas all other samples (which likely include some trees with full sibling and parent-offspring relationships) had genomic distances of between 0.051 and 0.235 (μ = 0.214). The latter included 13 pairs of trees with genomic distances less than 0.1, which were likely to be close relatives (Table S2). The genomic distance between our technical replicates suggests that we can expect 5-9% of genotypes to be miscalled, likely due to low sequence coverage. For technically replicated samples, the sequence set with the highest coverage per replicate pair was used in all subsequent analyses.

#### Genomic Prediction of Susceptibility

A genetic estimated breeding value (GEBV) for each tree was calculated using effect sizes obtained from field trial data as described in Stocks et al. 2019(*26*). Briefly, the trials comprised ash trees from 10 UK native seed zone (NSZ) provenances -including the zone in which Marden Park is found -together with one from Ireland and one from Germany. The trial trees were of similar age (∼7 years) planted in randomized plots, with even spacing, on former agricultural land. For the genome-wide association study (GWAS) 38,784 trees were scored in the trials for ash dieback damage using a health assessment method very similar to the one we used on the juvenile trees in Marden Park wood. Only 792 fully healthy trees were found in the trials (1.96% of all trees), and of these 623 were used for GWAS and to train the GP model, together with 627 highly damaged trees, giving a total of 1250 trees selected from the extremes of ash dieback damage phenotypes (*26*). An rrBLUP model was trained using pool-seq data from these 1250 trees, based on 10,000 loci that had the most significant association with disease susceptibility in a pool-seq GWAS (*26*). These 10,000 gave GEBVs that were able to predict the disease classification of a test population of 148 individually sequenced test trees with 68% accuracy (*26*).

The proportion of phenotypic variation explained by these GEBVs (i.e. the SNP heritability) was not calculated in (*26*). Hence, to now estimate the proportion of phenotypic variation in the latent variable explained by the GEBVs estimate, we conducted an abc analysis using the abc package in R (*50*), using the loclinear method with a tolerance of 0.1. The test statistic was the difference between the GEBV of the diseased and healthy plants in the test population of (*26*), divided by the standard deviation. This value was modeled using random draws from a normal distribution with the same sample size as the real data, and the same proportion of the total population in the diseased and healthy categories in (*26*) as measured in (*51*). The estimate of the proportion of genetic variation measured by GEBV was estimated by the posterior distribution of a parameter h, which was given a uniform prior density between 0.1 and 0.4.

In our data from Marden Park wood, 7,985 of these 10,000 “predictive” SNPs were variable within the population. The 7,985 SNPs were distributed among 2,783 contigs within our reference genome, which has contig N50 of 25kb. Only 32% of GEBV SNPs have another within 25 kb on the same contig, at which distance the average LD is below r²=0.08 (*45*). Effect sizes from the Stocks et al model, characterized as the additive effect of each copy of the alternate allele, were used to calculate GEBV of the Marden Park trees as *GEBV_j_* = Σ(*g_j_ f_ij_*) + 1, where GEBV_j_ is the genomic estimated breeding value of individual j, *g_j_* is the effect size at site i (Supp. Data 2 (*52*)), and *f_ij_ <u>ɛ</u>*{0,1,2} is the frequency of the alternate alleles at site i in individual j (Supp. Data 3 (*52*)).

We assessed the relationship between GEBVs and tree health phenotypes by correlation of trees’ GEBVs with the health scores of juvenile trees and the percentage canopy cover of the adult trees (Fig. S3). The mean difference in health phenotypes of the trees in equal numbers of trees with the highest and lowest GEBVs were compared in increments of one tree and plotted in terms of percentage of trees included (Fig. S4), finding a weak trend in the expected direction, which was not formally significant.

We also assessed the relationship between GEBVs and a second measure of tree health: the rate of increase in crown greenness as the adult trees flushed in early summer. The change was surveyed by an unmanned aerial vehicle (see above). The rate of crown greening was estimated by fitting a sigmoid function to the time series data of the normalized green chromatic coordinate:

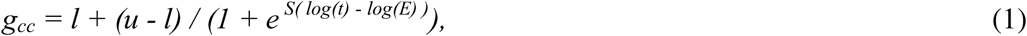

using the non-linear option of the R package brms (*55*),(*56*). This equation describes a curve between lower and upper values (*l* and *u*), with a slope, *S*, and mid-point (*E*).

The model was:

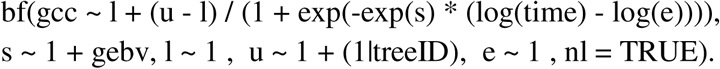

In this form convergence has been improved by parameterising the slope parameter as S, where *S* = *e^s^*. Time was standardized to the range [0,1] (13). The values of *u* were considered specific to each tree, and therefore modeled as a random effect, u ∼ 1 + (1|treeID). The rate of greening was modeled as a linear function of the GEBV score, s ∼ 1 + gebv. The gebv coefficient showed moderate evidence of a positive effect (the 95% CI was positive, Table S4).

#### Assignment of parentage

Parentage assignment was performed with the sequoia R package (V2.3.1) (*35*), using a randomly selected set of 1,000 SNPs having read depth > 20, minor allele frequency > 0.4, and an estimated error rate < 0.01. Parentage assignment was run using the hermaphrodite “B” mode. DBH was used to create a proxy for the birth year of adult trees (birth year = 100 -DBH) and juvenile trees were assigned a birth year of 100. This allowed for larger, and therefore presumably older, adult trees to be assigned as parents of younger adult trees, while disallowing juveniles from being considered as parent trees. The maximum age of parents was set as 99, allowing all adult trees to be considered as parents of the juveniles. The proxy years were not intended to be an accurate estimate of tree age. Candidate relationships were assigned based on a log likelihood threshold of −2. Overall confidence of parentage assignment was estimated within sequoia by simulating genotype data based on the estimated pedigree, recalculating parentage assignment based upon simulated data and comparing the recalculated pedigree to the original pedigree. The proportion of correct assignments over multiple runs of the process gives an estimate of confidence in our parentage assignments.

Parentage was assigned to 153 trees, with an overall confidence value of 0.72, obtained by repeatedly simulating genotype data from a reference pedigree, reconstructing that pedigree and counting accurate assignments. For 125 trees we could assign a single parent and for 28 trees we could assign two parents. Of trees with assigned parentage, 121 were classified as juveniles and 16 as adults. We identified 36 adult trees as likely parents, with a mean of 4.9 offspring per parent tree. Twelve of the trees were only assigned one likely offspring and 18 trees were assigned at least three offspring (Table S3). The largest set of offspring (n = 29) came from tree 51, which was both large and centrally located (Fig. S1), accounting for 21% of all trees with assigned parentage. Of these 29 offspring, seven had tree 10 (located in close proximity to tree 51) as the other parent, making them full siblings.

To test if parental GEBV was a better predictor of offspring health than parental phenotype, GEBV and percentage canopy cover of parent trees assigned by sequoia were each fitted against the mean health score of their likely offspring in linear models, weighted according to the number of likely offspring assigned to each parent tree (Fig. 2). The fit of each model was summarized as the Pearson’s correlation coefficient.

#### Detecting shifts in breeding value between adult and juvenile trees

We wished to investigate whether juvenile trees had higher GEBVs than we would have expected, based on the GEBVs of their adult relatives. All else being equal we would expect closely related individuals to have similar GEBV values. We should therefore be able to predict the GEBV of a tree from a sufficient number of SNPs that are unlinked to the GEBV sites and to each other, but which capture the pattern of relatedness. We therefore selected 5,793 such loci with a minor allele frequency of at least 0.3 (to ensure they were informative). To test whether these 5,793 loci were sufficient to estimate GEBV on the basis of relatedness, we divided the juvenile population into training and test groups with a ratio of 60:40, giving a training set of 272 juvenile trees and a test set of 181 juvenile trees. The split of the data was repeated 100 times and an rrBLUP(V4.6.1)(*57*) model was fitted on the training set as: *GEBV_J_* = *µ* + *x_J_β_J_* + *ε*, where GEBV_j_ is the vector of breeding values of juvenile trees, µ is the mean, *X_J_* is a matrix of juvenile genotypes (∈ {0,1,2}, giving the number of copies of the alternate allele) for the 5,793 SNPs, *β_J_* is a vector of allelic effects (treated as normally distributed random effects) and the residual variance is *Var*[ε]. Effect sizes from this model were then used to calculate rGEBV for the 181 tests trees as: *rGEBV_J_* = *µ* + *Σ*(*g_j_ f_ij_*), where *g_j_* the effect size at site i, and f_ij_ is the genotype at site i in individual j. For each of the 100 runs, the correlation of previously estimated GEBV and rGEBV was tested using a Peason’s product moment correlation and found to be significant (p < 0.01) in all 100 runs with a mean r = 0.50 (sd = 0.04). The correlation of rGEBV with GEBV demonstrates that this selection of SNPs are sufficient to predict GEBV based on relatedness. We then used the 5793 SNPs, to calculate an additive related matrix, which was used to predict the GEBV of juveniles from their relatedness to adults, which we then compared to their GEBVs based on the 7985 loci associated with ash dieback resistance. The t-statistic of the intercept was computed by the lm function of R, and evaluated against the t-distribution with 452 degrees of freedom.

#### Comparison of shifts in GEBV and unlinked loci

The variance in allele frequencies between adults and juveniles was quantified by the statistic F =∑ 2(Δp)^2^ / N (where Δp is change in allele frequency between generations and N the number of loci). These values were calculated for the set of 5793 unlinked SNPs (F_u_) and for the 7985 predictive loci (F_g_). Because the sampling distribution will depend on the pattern of linkage disequilibrium and the allele frequency spectrum, it was calculated empirically by randomizing the allocation of individuals to adult and juvenile categories, and the F estimates adjusted by subtracting the mean. The P-value for the difference *D* = F_g_ - F_u_ was the empirical cumulative distribution (ecd) at *D* in the null distribution (generated as differences between pairs drawn from empirical distributions above). To partition the variance of the 7985 GEBV loci into the contribution of selection and genetic drift, we modelled the selective component as a quadratic function of s = *p(1-p),* where *E* is the estimated effect size and *p* the allele frequency at each locus. This quadratic curve was used to calculate the contribution of drift (the F values in the absence of selection, i.e. at *s* = 0). The P-value was calculated as the ecd of the difference in the null distribution calculated in the same way as the unadjusted values.

#### Allelic shifts at causative loci

If allele shifts are due to natural selection we would expect the magnitude of shifts to be related to the effect size of the allele and the frequency of the allele in the population. The expected change in allele frequency under selection, from *f* to *f’,* is given by the standard equation *f’ =* (*f^2^ + f(1-f)(1-s))/w,* where s is the selective effect and w the mean fitness.

The value, *Δf* = *f’-f,* is therefore proportional to *sf*(1-*f*). Under the assumption that the selective effect was proportional to the estimated effect sizes *g_A_*, we plotted the observed change in minor allele frequency, *Δf*, against *g_A_f*(1-*f*) in Fig. 4. To visualize the trend more clearly we grouped the loci into 200 bins (quantiles of *g_A_f*(1-*f*)). The means of each bin are plotted in with an area of each point being inversely proportional to the variance in *Δf*, to convey the relative precision of the mean*Δf*, estimate. The line is the fitted linear regression (carried out on the individual Llf values) and the area of each symbol is proportional to 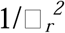 showing the relative weight attributable to each point.To test whether the regression was affected by the precision of the estimates of *E* for each locus (as measured by its standard error *SE_i_*) the regression was repeated with each locus inversely weighed by 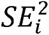.

#### Prediction on 121 juveniles with known parentage

For trees with assigned parents the expected frequency of the minor allele was calculated as; 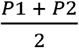, where *P*1 was the frequency of the allele in parent one and *P*2 was the frequency in parent two. Where only one parent was identified the mean frequency of the minor allele across the whole of the population was used as *P*2. The 7,985 sites of interest were tested for significant deviation between expected and observed allele frequencies on a per site basis using a two sided Wilcoxon rank sum test with continuity correction, with p-values corrected for multiple testing using the Bonferroni correction. Correlation between effect size and the mean difference between observed and expected allele frequency in offspring at each site was tested using Pearson’s product-moment correlation.

#### Exploration of polygenesis

We calculated the proportion of total variance in GEBVs contributed by the loci on each 1 kbp segment of our contig assembly, and how many of these were needed to explain 90% of the variance. We re-ran the analysis of shift in GEBV between adults and juveniles excluding the segments of the genome accounting for the top 25% of variance.

#### Estimating error on effect size estimates using simulated data

A simulation of the study was carried out to evaluate the effectiveness of the study design. The simulation consisted of two stages.

Firstly, we simulated the pool-seq genomic prediction field experiment. An initial population of 2983 simulated trees were allocated a genotype at 7985 loci by random binomial draw using our empirical estimates of the distribution of allele frequencies in the Marden Park population. Breeding values were calculated for these trees as *BV* = *X_J_β_J_*, where β*_J_* is the empirical estimate of effect sizes obtained in Stocks et al. (2019) for each locus and *x_J_* is the matrix of allele frequencies. We used the variance of the breeding values of these trees to calculate the size of a random environmental effect required for environment to be responsible for 60% of phenotypic variation (i.e. confer a heritability of h = 0.4). The environmental variance (*V_e_*) was calculated from heritability as 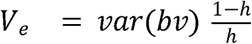. The underlying latent variable describing the susceptibility phenotype of all 2983 trees was generated by adding the random environmental effect to the breeding value: 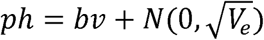.

In the pool-seq genomic prediction field experiment, 1300 out of 38784 phenotyped trees were included in the pools: 650 of the most healthy trees and 650 damaged trees. We therefore took a similar proportion of trees in the simulated population of 2983 trees to make two pools: the 50 highest phenotype values and the 50 lowest. This enabled us to define the upper and lower quantiles of the latent variable describing the susceptibility phenotype. For efficiency of simulation, to produce 30 pools of 40 trees, we repeatedly simulated new genotypes and calculated their phenotypes, retaining in pools those individuals with phenotypes lying outside of the upper and lower quantiles. These genotypes were added to the 15 upper and 15 lower pools until each contained 40 individuals. We then analyzed these pools in a similar way as the pools from the field trial in Stocks et al. (2019). Using the rrBLUP mixed.solve algorithm we estimated effect sizes for the 7985 sites, from their allele frequencies in the 15 upper and 15 lower pools.

We then compared these estimated effect sizes with the actual effect sizes that each locus had been given at the start of the simulation (from Stocks et al 2019), by correlation.

---

**Fig. S1.**
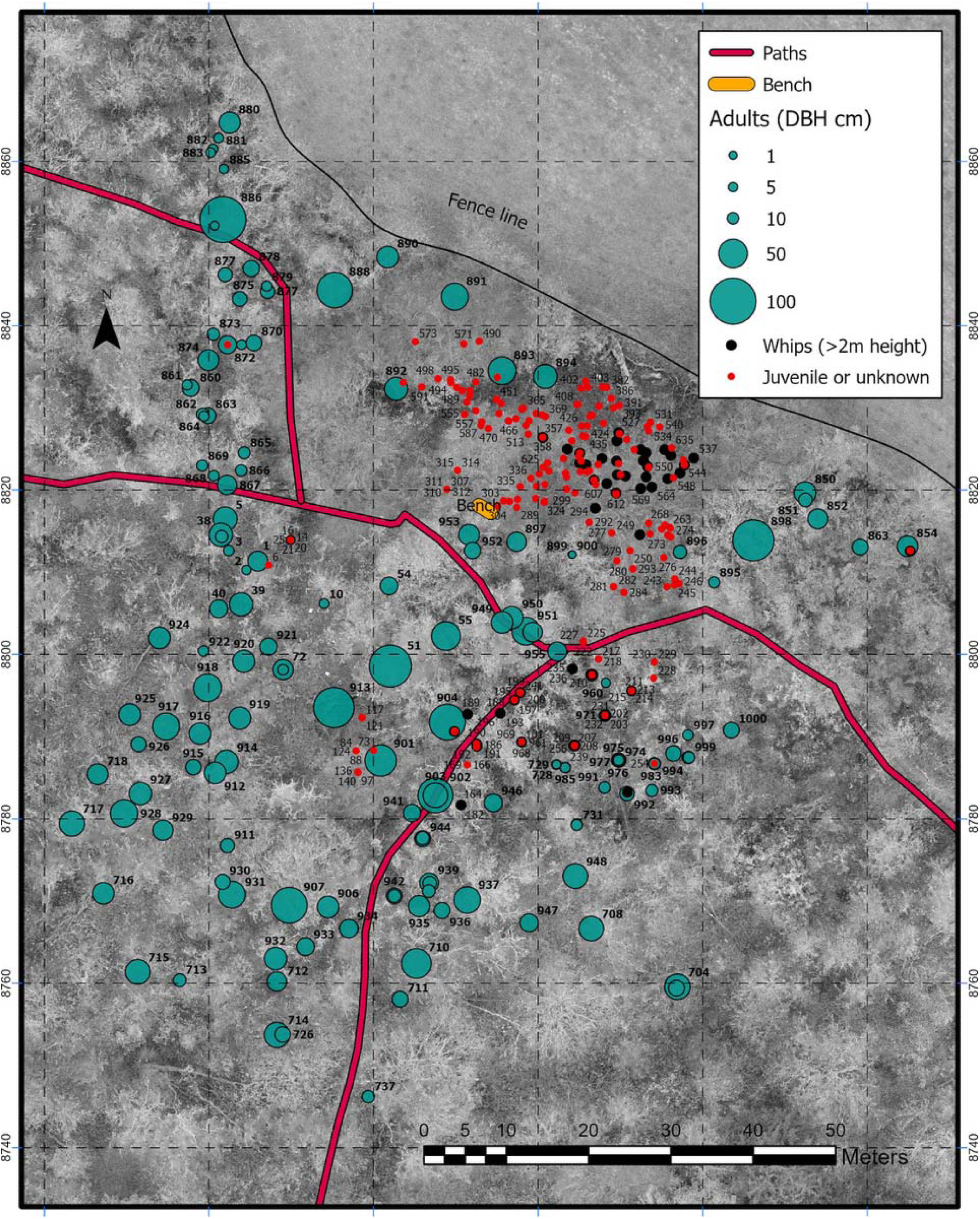
Map of samples showing location of trees at the Marden Park site. Collected in March 2021 using an RTK GPS (Bad Elf Pro), using NTRIP (Networked Transport of RTCM via Internet Protocol) connection to the Imperial University unit (LICCOOGBRO), approximately 27 km from the site. Background image from UAV campaign February 2022.

**Fig. S2.**
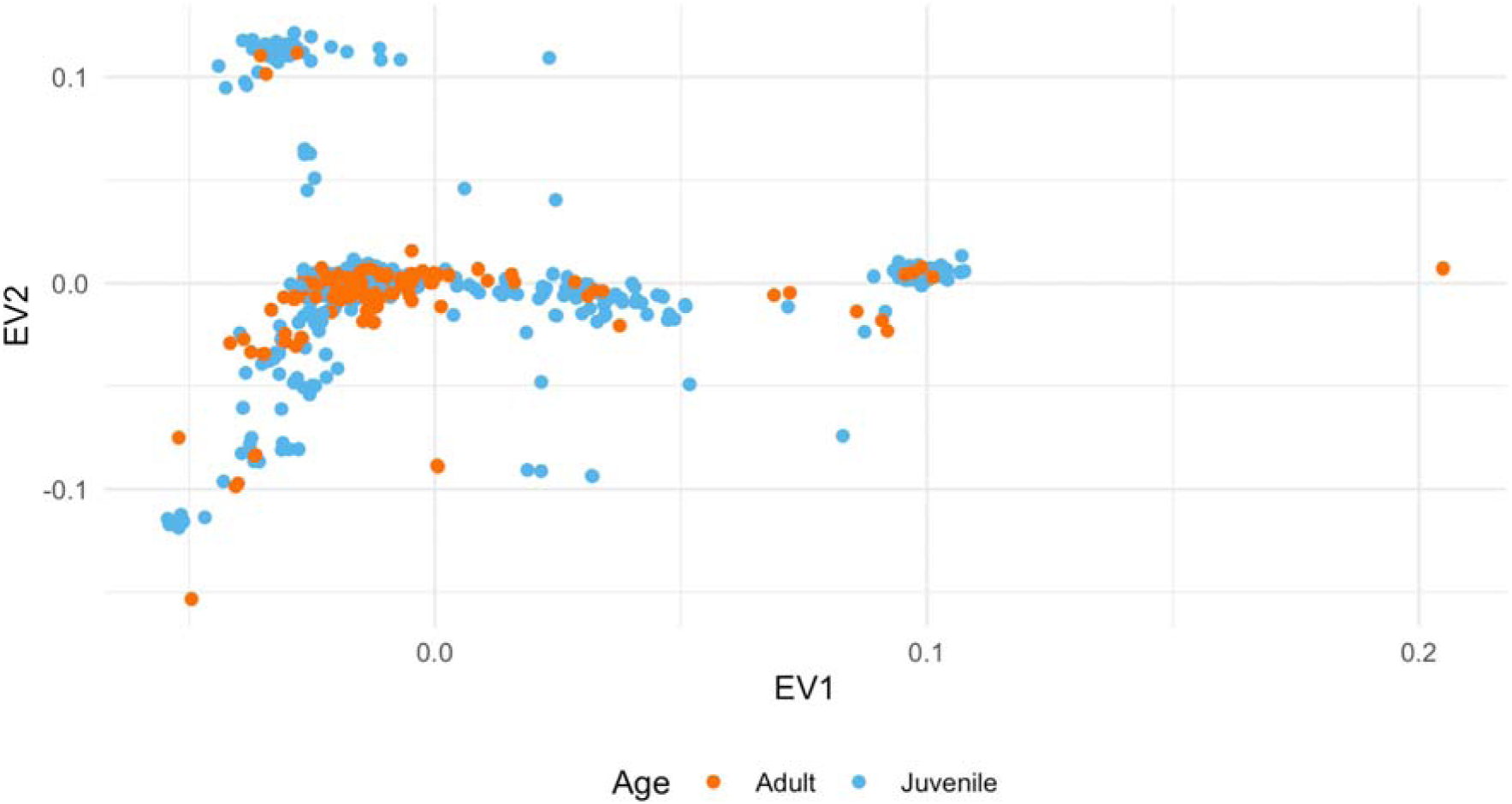
Principal genetic components of all adult and juvenile ash trees, including technical replicates, based on 4,287,210 SNPs.

**Fig. S3.**
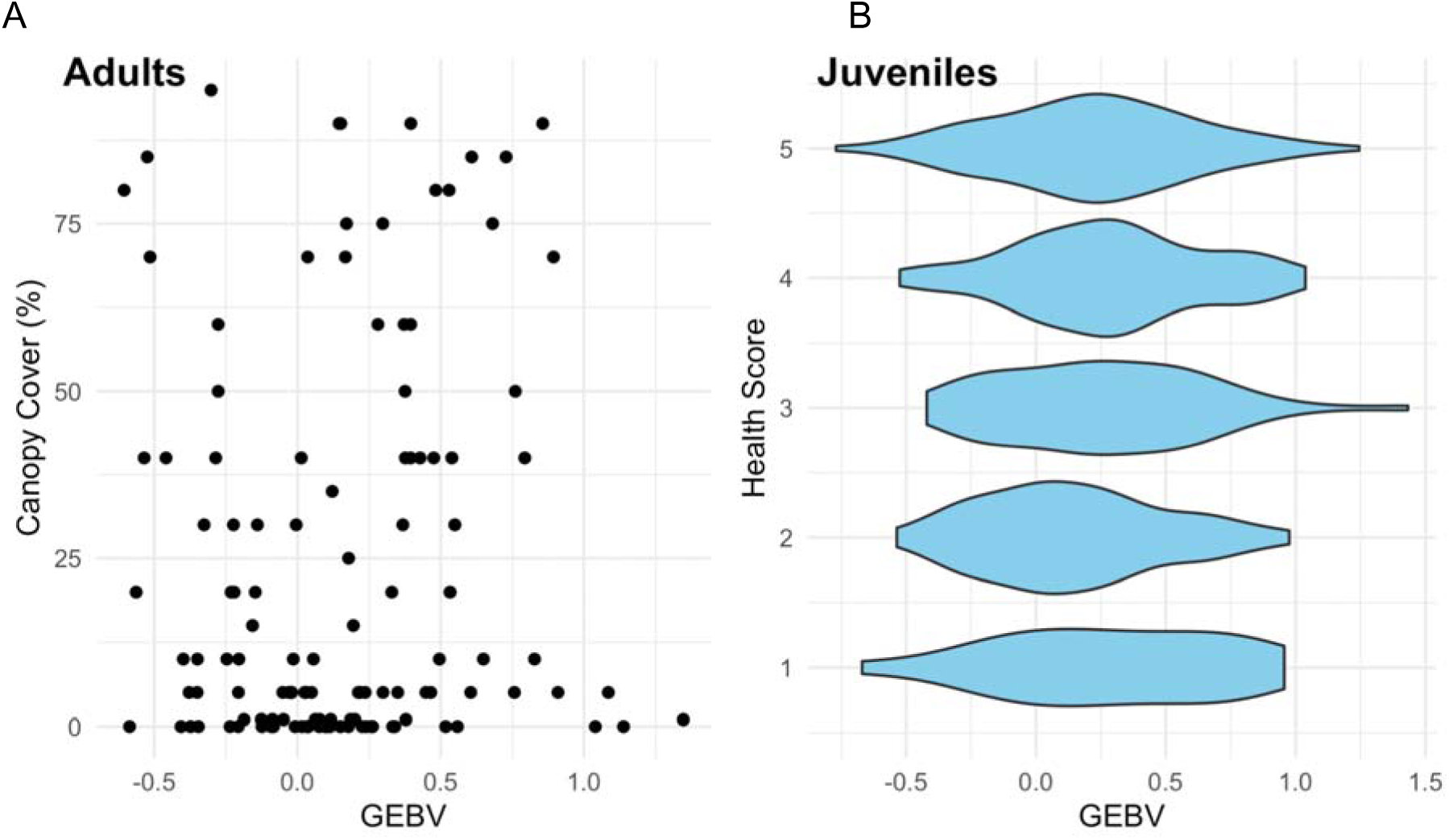
Relationship between genomically estimated breeding values (GEBV) and phenotypic estimates of ash dieback damage. (A) GEBV of 128 adult trees using 7,985 loci plotted against percentage canopy cover as scored in 2019. (B) GEBV of 452 juvenile trees using 7,985 loci plotted against 2019 phenotypic scores on a 1-5 scale where 1 is most severely affected.

**Fig. S4.**
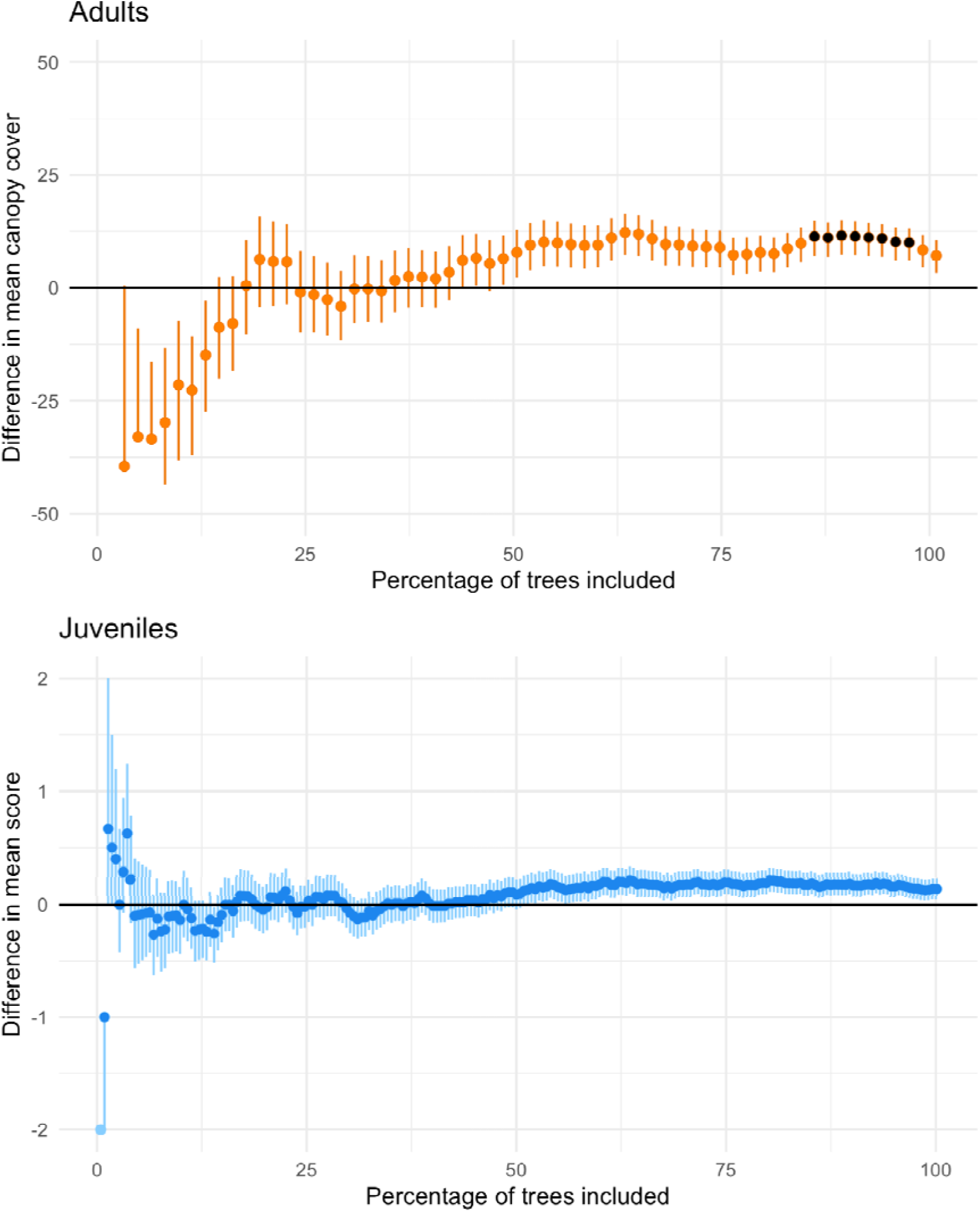
Difference in phenotypic scores of ash dieback damage for trees in equally sized upper and lower extremes of the GEBV distribution. (A) Percentage canopy cover of adult trees (n = 128) and (B) Health scores of juvenile trees (n = 452). GEBVs were calculated from 7,985 SNP loci. Bars represent the standard error on the cumulative mean phenotypes and points in black show a significant difference (paired-t test, p <0.05) in mean phenotype between the trees in the upper and lower extremes of the GEBV distribution.

**Fig. S5.**
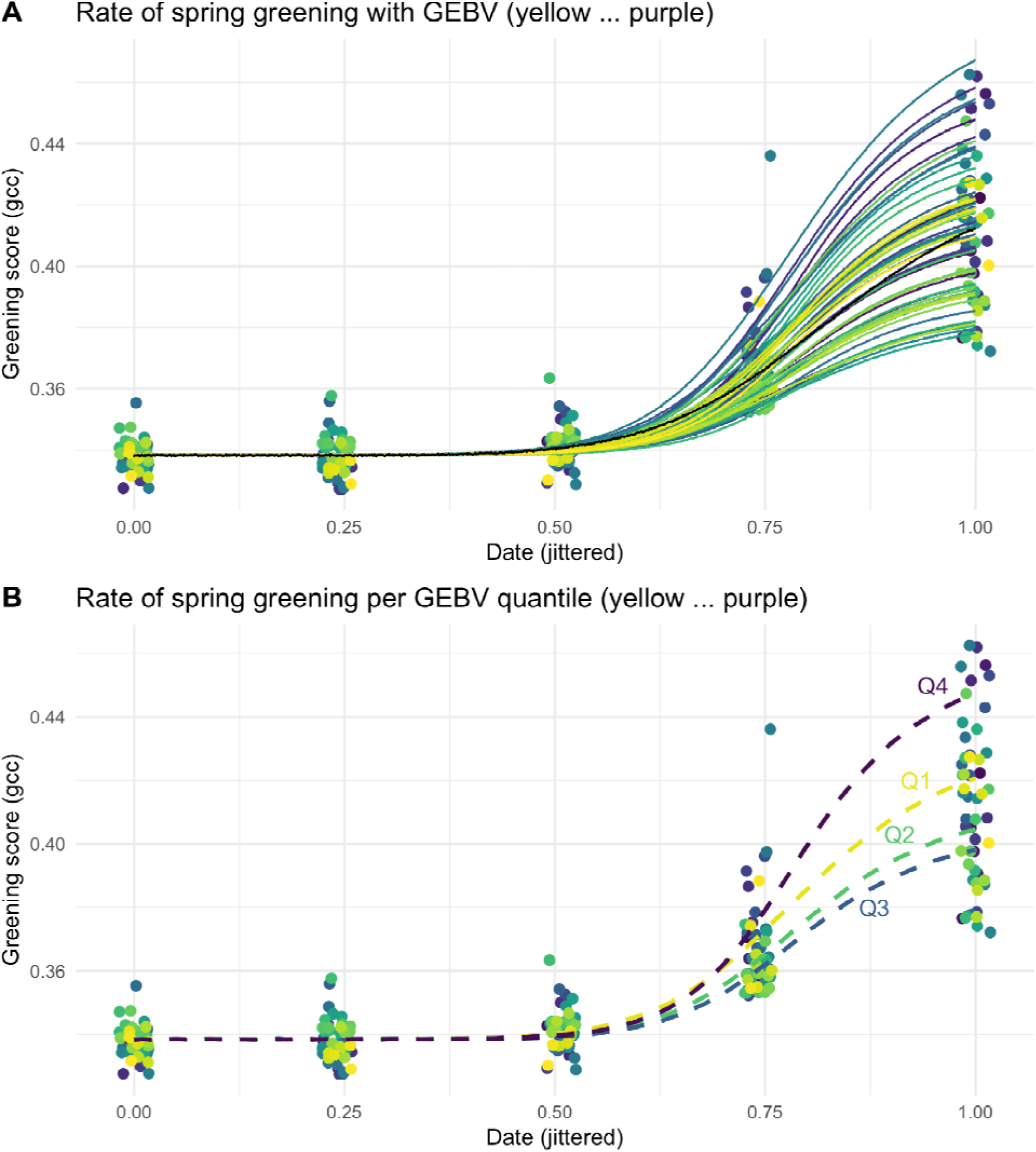
The rate of spring green-up as a function of GEBV. The points are the observed values recorded by the UAV (drone) for each tree. A) The lines are the fitted values for each tree. The colors indicate the relative GEBV score (ranging low to high from yellow to purple). The trees with a higher GEBV score have a faster rate of greening. B) Lines fitted for each quartile.

**Fig. S6.**
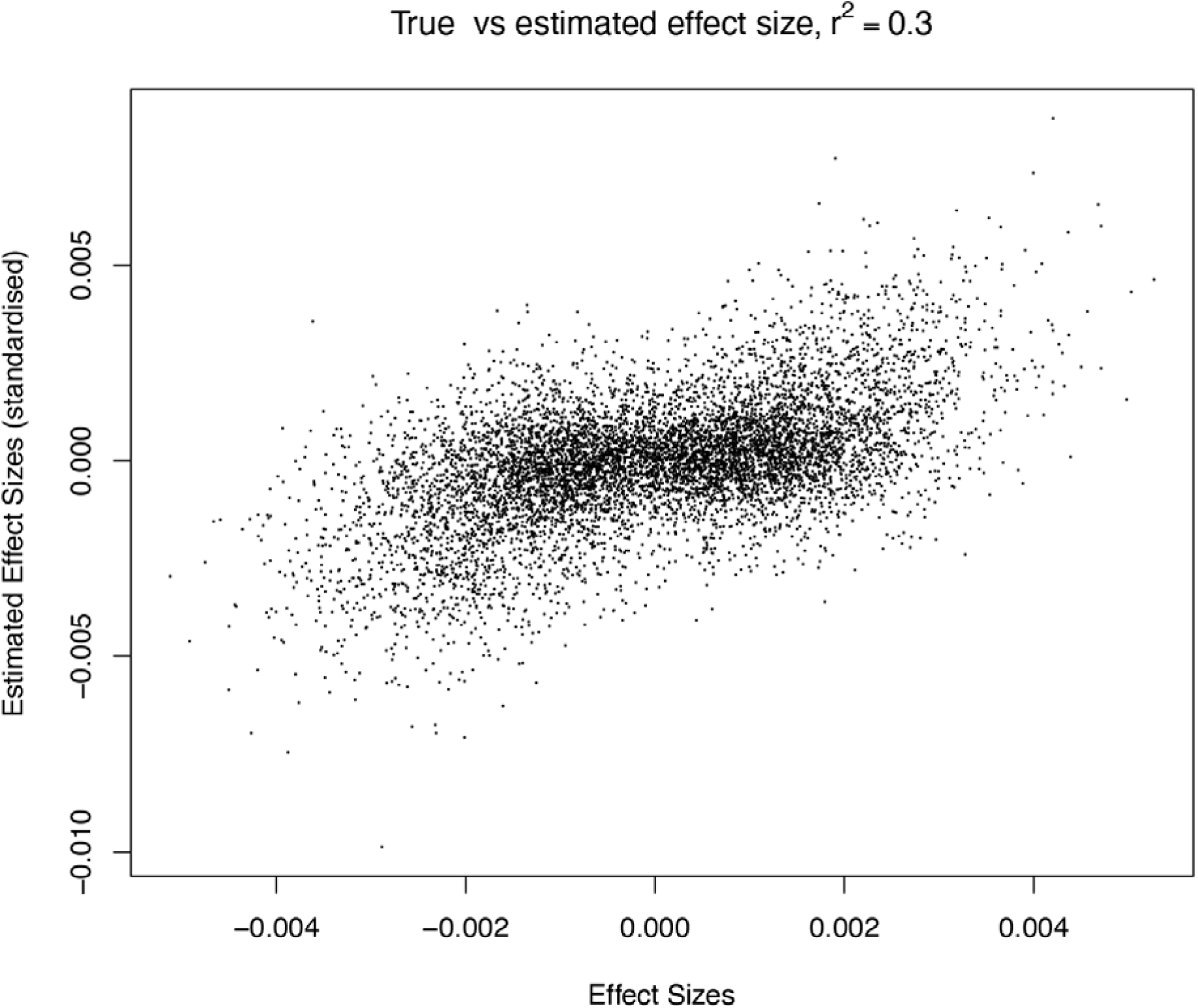
True versus estimated effect sizes of 7985 loci in a simulation of our experimental design. Effect sizes measured in Stocks et al (2019) were used as the true values. These values were used to calculate the phenotypes of a set of trees, simulated with random genotypes and a random environmental effect that conferred a heritability of 0.4. These phenotypes were used to train a genomic prediction model estimating the effect size of each locus. The relative effect sizes were estimated convincingly, albeit with some error (r^2^=0.30).

**Fig. S7.**
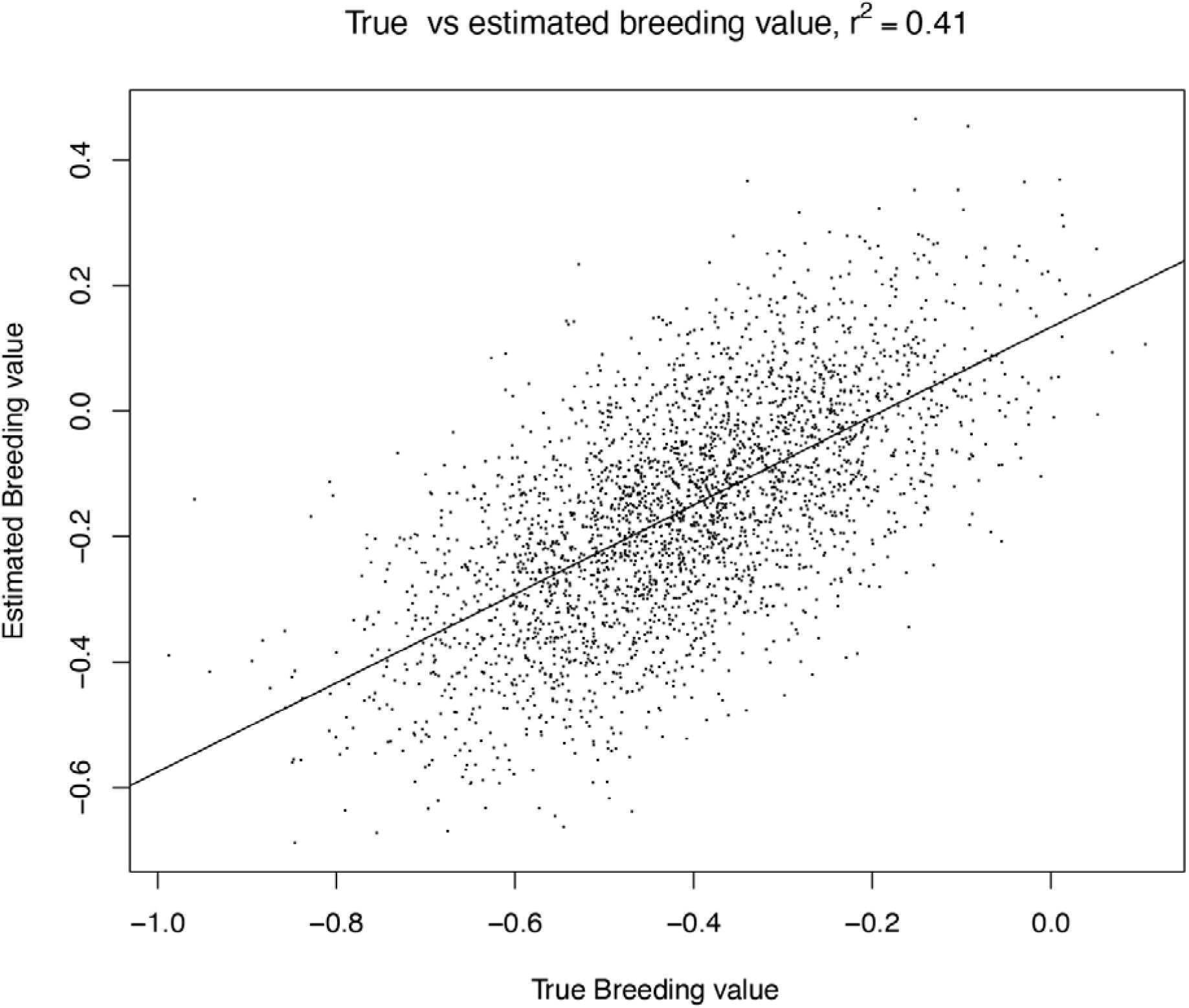
True breeding value versus estimated breeding value of simulated trees. True breeding values were calculated from the genotypes of the simulated trees using the effect sizes for each locus from Stocks et al (2019). Estimated breeding values were calculated by summing the effect size estimates shown in Fig. S7 for each simulated tree. Even though the effect size of each locus was estimated with considerable error (Fig. S7), when the effects at multiple loci are summed, they allow for accurate genomic prediction (r^2^=0.41).

**Fig. S8.**
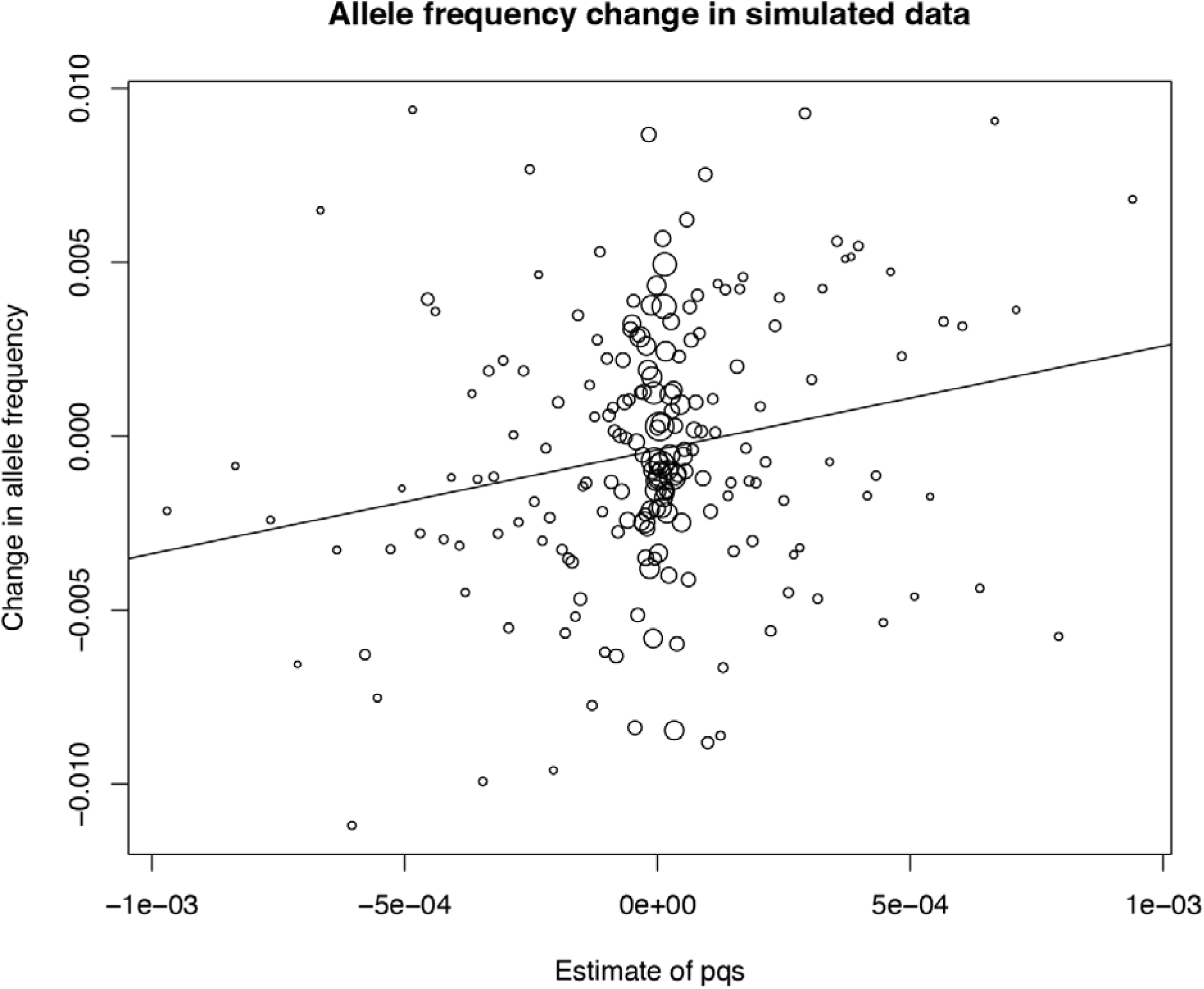
Simulation of the Marden Park population showed shifts in allele frequencies that correlated with the effect size and allele frequency of each locus (multiple r^2^ = 0.0011, p = 0.002452, F = 9.102 on 1 and 7983 DF), similar to that found in our field study (see Fig. 3).

**Table S1.**
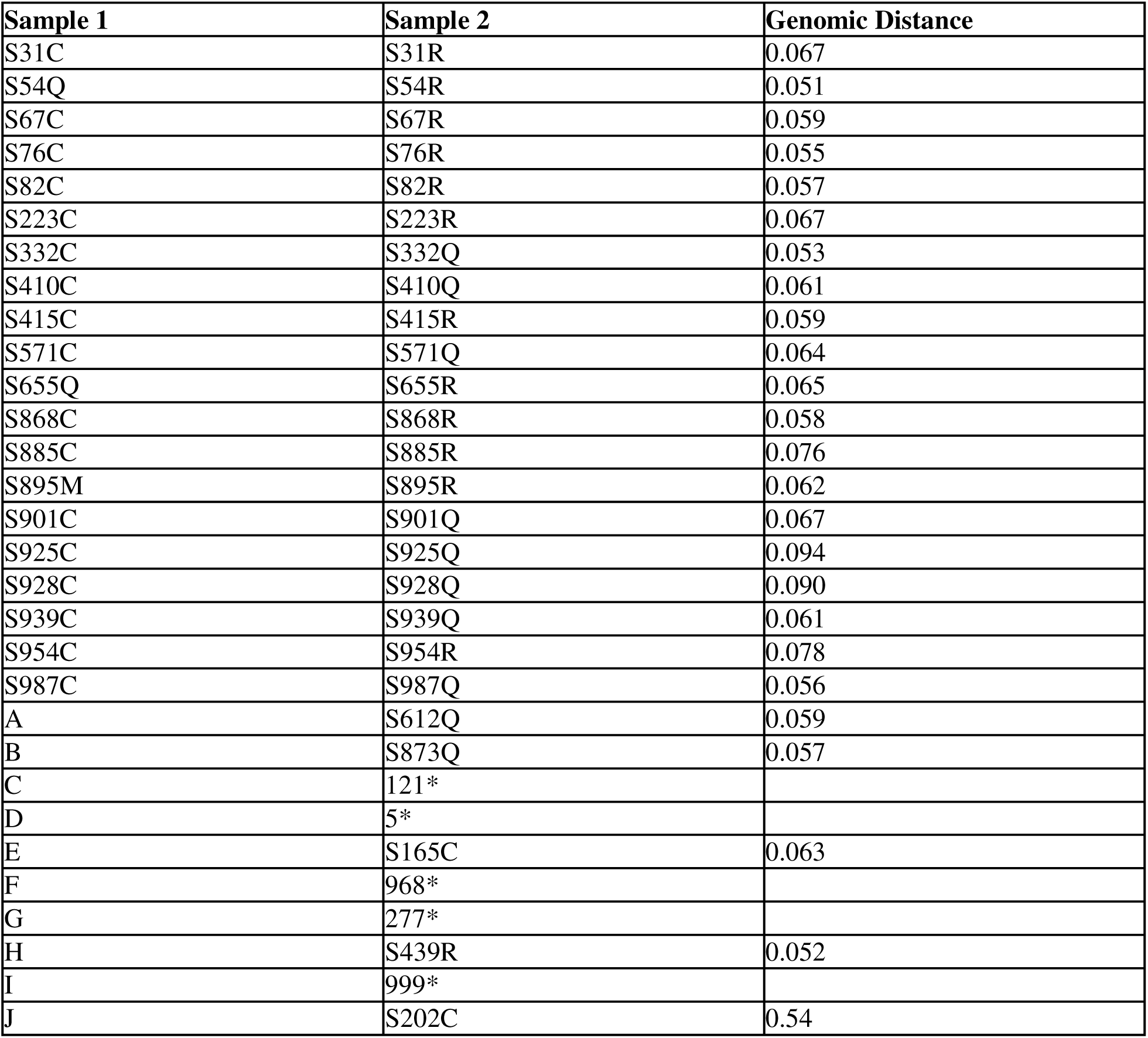
Technical replicate identification. Genomic distance (1 minus identity-by-state) between replicate samples. For blind replicates C, D, F, G and I, their duplicate did not pass read alignment thresholds and was therefore not included in SNP calling. In these cases the replicate was relabelled to act as a replacement sample.

| Sample 1 | Sample 2 | Genomic Distance |
| --- | --- | --- |
| S31C | S31R | 0.067 |
| S54Q | S54R | 0.051 |
| S67C | S67R | 0.059 |
| S76C | S76R | 0.055 |
| S82C | S82R | 0.057 |
| S223C | S223R | 0.067 |
| S332C | S332Q | 0.053 |
| S410C | S410Q | 0.061 |
| S415C | S415R | 0.059 |
| S571C | S571Q | 0.064 |
| S655Q | S655R | 0.065 |
| S868C | S868R | 0.058 |
| S885C | S885R | 0.076 |
| S895M | S895R | 0.062 |
| S901C | S901Q | 0.067 |
| S925C | S925Q | 0.094 |
| S928C | S928Q | 0.090 |
| S939C | S939Q | 0.061 |
| S954C | S954R | 0.078 |
| S987C | S987Q | 0.056 |
| A | S612Q | 0.059 |
| B | S873Q | 0.057 |
| C | 121* |  |
| D | 5* |  |
| E | S165C | 0.063 |
| F | 968* |  |
| G | 277* |  |
| H | S439R | 0.052 |
| I | 999* |  |
| J | S202C | 0.54 |

**Table S2.** Potential Close Relatives among our samples. Genomic distance (1 minus identity-by-state) between similar samples not labeled as replicates.

| Sample 1 | Sample 2 | Genomic Distance |
| --- | --- | --- |
| 7 | 17 | 0.074 |
| 16 | 110 | 0.059 |
| 44 | 951 | 0.053 |
| 156 | 160 | 0.061 |
| 207 | 209 | 0.069 |
| 281 | 282 | 0.074 |
| 281 | 283 | 0.073 |
| 282 | 283 | 0.065 |
| 458 | 461 | 0.058 |
| 470 | 560 | 0.069 |
| 472 | 877 | 0.063 |
| 473 | 476 | 0.064 |
| 533 | 549 | 0.064 |
| 903 | 906 | 0.064 |

**Table S3.** Parentage assignments. Showing the number and mean (where number of offspring > 1) health scores of offspring assigned to each likely parent tree. An accurate DBH measurement is missing for tree 10.

| Parent Tree | DBH of parent (cm) | % canopy cover in parent (2019) | Parental GEBV | Number of likely offspring | Mean score of offspring (2019) |
| --- | --- | --- | --- | --- | --- |
| 51 | 80.0 | 90 | 0.8720 | 29 | 4.3 |
| 10 | - | 5 | 0.0087 | 20 | 4.5 |
| 889 | 42.2 | 5 | -0.3587 | 19 | 3.9 |
| 39 | 37.0 | 75 | 0.1908 | 15 | 3.5 |
| 71 | 49.8 | 5 | 0.2481 | 12 | 4.1 |
| 53 | 48.1 | 80 | 0.5806 | 8 | 3.7 |
| 890 | 33.0 | 10 | -0.3074 | 8 | 3.0 |
| 937 | 45.0 | 10 | -0.3026 | 7 | 2.8 |
| 853 | 39.3 | 30 | 0.3898 | 6 | 3.0 |
| 860 | 22.3 | 0 | 0.3718 | 5 | 4.0 |
| 945 | 28.3 | 60 | 0.3248 | 5 | 3.8 |
| 850 | 34.5 | 40 | -0.4849 | 4 | 2.3 |
| 898 | 78.4 | 85 | -0.5119 | 4 | 2.0 |
| 901 | 55.5 | 0 | -0.2281 | 4 | 5.0 |
| 872 | 24.7 | 5 | 0.7402 | 3 | 2.3 |
| 902 | 64.0 | 10 | 0.0427 | 3 | 4.0 |
| 907 | 66.0 | 80 | -0.5879 | 3 | 4.7 |
| 920 | 34.0 | 5 | -0.0741 | 3 | 5.0 |
| 38 | 37.5 | 5 | -0.3661 | 2 | 3 |
| 55 | 50.1 | 50 | 0.3625 | 2 | 4.5 |
| 884 | 89.8 | 70 | -0.3513 | 2 | 3 |
| 886 | 40.0 | 5 | 0.0052 | 2 | 2 |
| 908 | 4.5 | 0 | 0.5565 | 2 | 5 |
| 910 | 4.5 | 20 | 0.3279 | 2 | 3.5 |
| 52 | 35.0 | 5 | -0.2242 | 1 | NA |
| 712 | 28.0 | NA | 0.4506 | 1 | 5 |
| 852 | 29.6 | 10 | -0.2466 | 1 | 2 |
| 896 | 14.2 | 1 | 0.0870 | 1 | 2 |
| 903 | 40 | 0 | 0.2233 | 1 | NA |
| 927 | 35.0 | 1 | 0.4155 | 1 | 5 |
| 928 | 47.0 | 30 | -0.2549 | 1 | 2 |
| 934 | 26.0 | 5 | 0.1546 | 1 | 3 |
| 973 | 6.8 | 10 | 0.4924 | 1 | 5 |
| 978 | 13.2 | 0 | -0.3005 | 1 | 4 |
| 988 | 16.8 | 0 | -0.0603 | 1 | NA |
| 989 | 4.1 | 70 | 0.8080 | 1 | 5 |

**Table S4.**
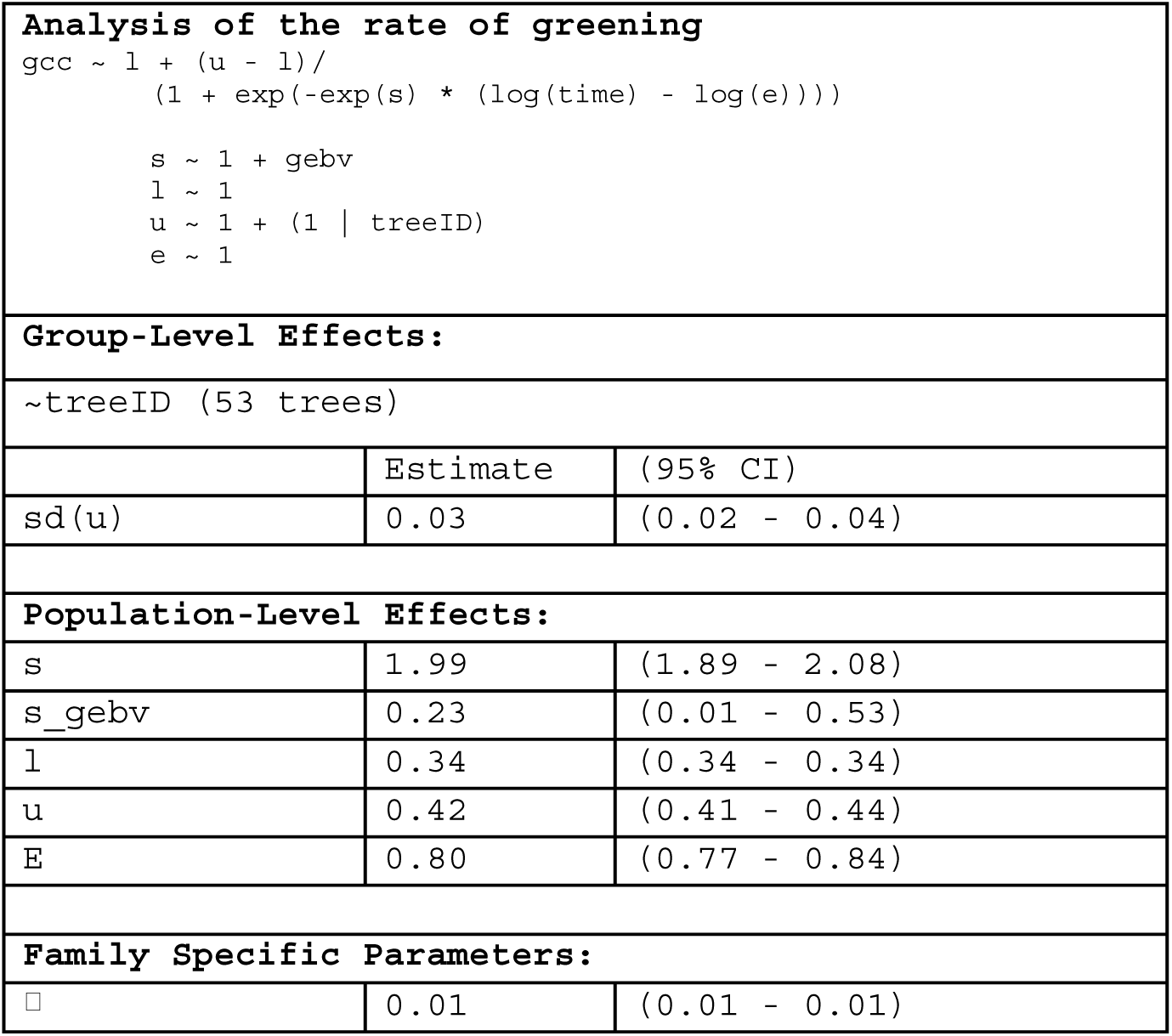
The fitted values of the model describing the early summer greening as a function of GEBV score.

